# Dissecting functional elements in giant genomes using the first generation of high-quality salamander assemblies

**DOI:** 10.64898/2026.10.27.762315

**Authors:** Nataliya Timoshevskaya, Vladimir A. Timoshevskiy, Kaan I. Eskut, Erich D. Jarvis, S. Randal Voss, Jeramiah J. Smith

## Abstract

Salamanders are an enduring and historically significant group that helped originate multiple scientific fields of study, including regeneration, cytogenetics, experimental embryology, cloning, evolutionary developmental biology and evolution of extreme genome size. We generated a high-quality, single haplotype assembly for the 32 Gb axolotl (*Ambystoma mexicanum*) genome using an F1 hybrid between axolotl and a divergent relative the tiger salamander (A. *tigrinum*). Using multi-genome alignments to other salamanders that meet VGP metrics, we leveraged strong conservation of karyotypic structure to identify >523,000 highly conserved non-coding elements within massively expanded intronic and intergenic regions, the vast majority of which are unique to the salamander lineage. Orthogonal measures of chromatin state and enhancer activity confirmed that many conserved elements correspond to cis-regulatory sequences and identified several thousand that are utilized during regeneration. The conserved elements identified here will enable robust functional genomic analyses of tissue regeneration and other traits that uniquely empower salamander models in biological research.

## INTRODUCTION

Tetrapod vertebrates present a diversity of shared and unique characteristics that reflect common ancestry and lineage specific evolution. At almost any comparable time of development, species within and among the major tetrapod groups show striking similarities and differences in structure and function that transcend all levels of biological organization. This is readily seen when comparing salamander characteristics to other amphibian, amniote, and mammalian taxa^1–5^. In comparison to other amphibians (frogs and caecilians), salamanders have repeatedly evolved non-metamorphic modes of development to exploit relatively permanent bodies of water for totally aquatic life histories^6^. While salamanders develop four limbs like other tetrapod species, the order in which digits are specified is uniquely preaxial^7,8^. Also, unlike other extant tetrapods, salamanders can regenerate their limbs into the adult phase of life^9^. Equally amazing, salamanders uniquely evolved large genomes that on average are approximately 10x larger than the human genome^10^. Expanded genomes may contain uniquely evolved sequences for regulating traits like metamorphosis and limb regeneration^11^, however a test of this hypothesis awaits comparative studies of salamander genomes.

As all living salamander species have large genomes, expansions in genome size likely occurred early in their ancestry and over a protracted period of time, between the divergence of ancestral frog and salamander lineages ∼250 MYA and the basal divergence within salamanders ∼175 MYA^12^. This genome expansion occurred without whole genome duplication, presumably resulting from the insertion of mobile elements into intronic and intergenic regions^13^. The deep ancestry of these repeat expansions and subsequent accumulation of mutations rendered most of the genome nonrepetitive as it relates to sequence identity^14^. This mode of expansion has left axolotl with a number of protein coding genes (UKY_AmexF1_1: N = 24,121) that is similar to that of human (GRCh38.p14: N = 20,078, T2T-CHM13v2.0: N = 20,071) and other diploid tetrapods^15–17^, but with regulatory elements that are dispersed over much broader distances within the genome^13,18,19^. Theoretically, this additional intronic and intergenic DNA might also provide a larger pallet of formerly functional transposable element sequences from which novel regulatory elements might evolve.

Among vertebrates, the distribution and evolution of regulatory elements is best understood in mammals. Extensive studies have used chromatin-oriented approaches (e.g. ATAC-seq) and content (DNA methylation, histone acetylation and methylation) to identify genomic regions that may regulate transcription. Additionally, conservation tracks that use alignments across multiple mammalian (and deeper vertebrate) species have revealed non-exonic regions that have accumulated mutations at a slower rate than expected under neutral drift. Comparison of epigenetic state-based (ATAC, H3K27ac, H3K4me) and conservation-based methods for identifying candidate enhancers has shown that a majority of chromatin-defined enhancers have arisen more recently, within mammalian sublineages^20–22^. These lineage-specific enhancers show higher rates of turnover and smaller impacts on transcription relative to more deeply conserved placental or mammalian enhancers^20–22^, highlighting the need to integrate measures of evolutionary constraint when evaluating enhancer function. Overall, the average half-life of an enhancer in a mammalian genome has been estimated to be ∼130 million years^22^, a timeframe that falls roughly between the estimated basal placental divergence (∼100 MYA) and the divergence of the ancestral placental/marsupial lineages (∼160 MY)^23^. As such, a relatively small subset of enhancers are found to be conserved at deeper levels of vertebrate phylogeny^24^. With this understanding of the tempo of enhancer evolution in mammals, we anticipate that only a small fraction of enhancers will be identifiable in sequence comparisons between deeply divergent vertebrate orders or amphibian Classes.

We reasoned that performing comparisons directly among salamander species will be necessary to identify many of the meaningful regulatory elements present in their genomes. These comparisons are particularly relevant given the deep ancestry of the salamander lineage, and the fact that many traits such as enhanced regeneration and maintenance of archetypical tetrapod morphology are unique to salamanders (contrasting the highly derived morphologies of frogs and caecilians). However, historically the large genome size of salamanders has prevented such analyses, due to the challenges in generating high-quality assemblies and alignments between large genomes. Recent improvements in sequencing and assembly algorithms using long reads and haplotype phasing, particularly by the Vertebrate Genomes Project (VGP) and affiliates have made it possible to generate highly contiguous assemblies for salamanders and other species with large genomes^15^. Improvements in alignment algorithms have also permitted the generation of multispecies alignments incorporating hundreds of species, although these still face challenges in scaling to large genomes, given that the computational resources needed for generating alignments that scale approximately quadratic with genome size^15,25^.

To enable the discovery of conserved elements in salamander genomes, we generated a haploid phased assembly for a model salamander (*Ambystoma mexicanum*: axolotl) using a F1 *A. mexicanum* X *A. tigrinum* hybrid. We then utilized a synteny guided approach to generate multi-salamander alignments, and performed cross validation of epigenetic state-based and conservation-based identification of candidate regulatory elements. With these resources and approaches, we identify and characterize regulatory elements that contribute to conserved aspects of salamander biology, including limb regeneration.

## RESULTS

### An F1 Hybrid Phased Assembly for axolotl

Sequencing was performed using an F1 hybrid between an axolotl (*A. mexicanum*) from the University of Kentucky *Ambystoma* Genetic Stock Center (AGSC) and a wild captured tiger salamander (*A. tigrinum*) from Clay Hill Memorial Forest, Campbellsville University, Kentucky (Figure 1). The AGSC stocks are the source material for nearly all research colonies that currently study axolotls today, including all genomes sequenced to date^18,19,26^. Animals from laboratory colonies of axolotl contain some ancestry from a historical introgression from another tiger salamander lineage^27^ but maintain gross morphology and the paedomorphic condition typical of wild axolotls. To generate an assembly, PacBio HiFi reads (N = 79,181,288) and Arima Hi-C linked read pairs (N = 3,338,422,583 150bp read pairs) were generated for the F1, and short reads were generated from its axolotl and tiger salamander parents to enable trio-based haplotype phasing. Axolotl and tiger salamander differ at ∼1 in every 80 bases^28^, thus providing sufficient polymorphism to facilitate robust haplotype phasing during assembly.

**Figure 1:**
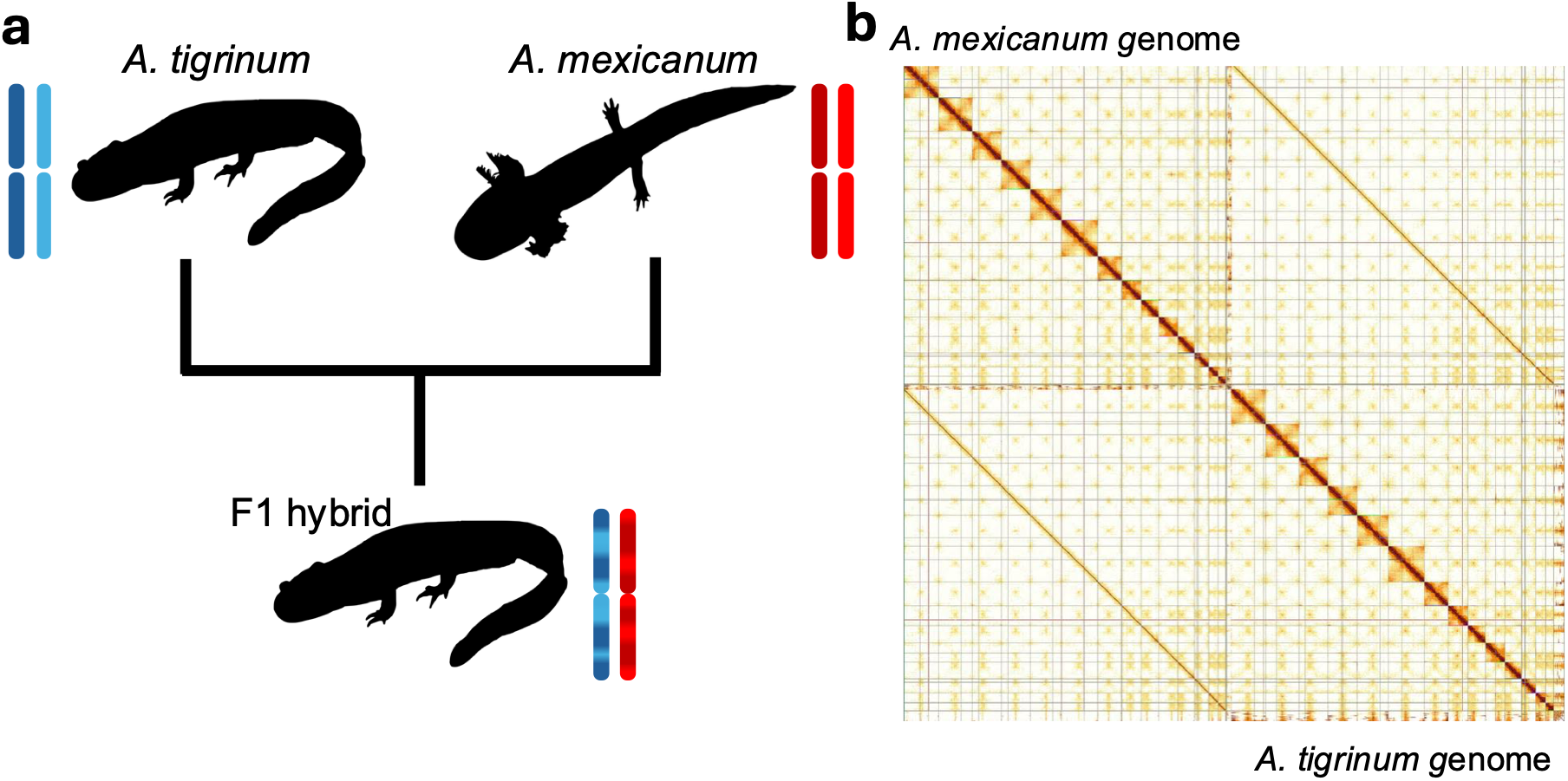
Strategy for assembly of *A. mexicanum* and *A. tigrinum* reference genomes. **a**) Interspecific crossing design used to generate haplotypic assemblies for both species. **b)** Hi-C contact maps generated for the F1 showing Hi-C contacts across all 14 axolotl and 14 tiger salamander chromosomes. All 14 chromosomes are resolved for both species. Vertical and horizontal lines intersecting individual chromosomes represent artificial breaks that were introduced into the seven largest chromosomes of each species to enable read mapping and a small number of manual scaffold curation events.

Assembly of HiFi reads using the HiFiasm assembly and paternal kmer profiles yielded separate assemblies for axolotl and tiger salamander haplotypes within the F1, with contig N50s of 23 and 26 Mb respectively, (*A. mexicanum*: N50 = 22,991,321, L50 = 387, N = 3618 contigs; *A. tigrinum* N50 = 25,538,197, L50 = 346, N = 4373 contigs) respectively. These assemblies were further scaffolded by aligning Hi-C data to both haplotype assemblies in the same alignment run. Subsequent dual manual curation was performed, considering alignments to both species simultaneously to ensure full chromosomal scaffolding and proper phasing (Figure 1). Scaffolds were oriented to ensure collinearity with previously published assemblies^18,19^, correcting two apparent inversion artefacts present in a recent draft^18^. The final axolotl assembly covered 29.1 Gb with 98.8% of the assembly assigned to chromosomes, the remainder largely corresponding to unlocalized rDNA and centromeric repeats. For tiger salamander, the assembly size (30 Gb) and percentage of unscaffolded sequence (3.7%) were slightly larger, presumably reflecting the additional impacts of known recent activity of long Harbinger elements in that genome^28^.

To account for current computational 32-bit limits of sequence size in NCBI submissions (2.1 Gb limit), the submitted sequences for largest chromosomes 1-7 were split at sequence gaps in the centromeric repeat arrays. The axolotl assembly was made publicly available under limited embargo of the Vertebrate Genomes Project in July 2024 to facilitate early use^29^ and was annotated by NCBI RefSeq using the NCBI Eukaryotic Genome Annotation Pipeline^30^. Associated BUSCO and proteome alignment analyses indicate a high degree of completeness (95.9% of tetrapoda_odb10 identified as complete), similar to or exceeding other amphibian RefSeq chromosomal assemblies^17^.

### Evolutionary conservation across salamander species

Analysis of the new axolotl genome assembly and available chromatin datasets validated expected patterns at proximate promoters and revealed millions of candidate cis-regulatory elements (CREs) (Supplementary Text, Figures S1-2). However, it should be recognized that epigenetic state-based methods are limited to the identification of CREs that accumulate signal in the specific cell types sampled. An alternate approach to regulatory element discovery is to identify regions of the genome that evolve slowly due to functional constraint. The accumulation of random mutations causes sequences to diverge from one another in regions of the genome that evolve at neutral or near neutral rates, however in regions where function is specified by the underlying sequence (e.g. protein coding regions or transcription factor binding sites) mutations are more likely to be selectively disadvantageous, causing these regions to diverge more slowly over time. As such, sequence conservation serves as a metric of sequence functionality over the deep evolutionary history of a lineage, reflecting millions of years of evolutionary experimentation across all genes, tissues and cell types. This core idea and the associated conservation tracks have been invaluable resources for studies of mammalian gene regulation for decades^31–34^.

To identify evolutionarily constrained sequences in the axolotl genome, we generated a reference free alignment of five available salamander genomes from sister projects that aim to produce high-quality genomes: axolotl (this study), *Pleurodeles waltl*^29^, *Triturus cristatus* (Darwin Tree of Life project, DToL)^35^, *Lissotriton helveticus* (DToL), and *Lissotriton vulgaris* (DToL). The later four species are members of the newt lineage (Family Salamandridae), which diverged from its common ancestor with *Ambystoma* ∼140 MYA. Altogether these five species sample ∼400 million years of salamander evolution, accounting for the length of all shared and terminal branches.

To circumvent scaling issues associated with alignment of large genomes, we took advantage of the relatively slow rate of karyotypic evolution in these two lineages^36^ by first using rapid pairwise alignments (minimap2)^37^ to identify groups of chromosomes that were associated by fission/fusion events. Using this strategy, we were able to define 10 evolutionarily independent subsets of chromosomes tracing to the common ancestor of newts and ambystomatids (Figure 2, Figure S3, Supplemental Table 1). These subsets of chromosomes were then aligned using Cactus reference-free aligner^25^, which ran to completion in approximately 60 days using two compute nodes with 128 processors and 5 TB of RAM each. After initial examination we identified one additional complex syntenic relationship that required an 11th alignment run to cover the full length of axolotl chr 3. The resulting alignment files were used to generate phyloP^33^ conservation tracks relative to the axolotl genome (Figure 3).

**Figure 2:**
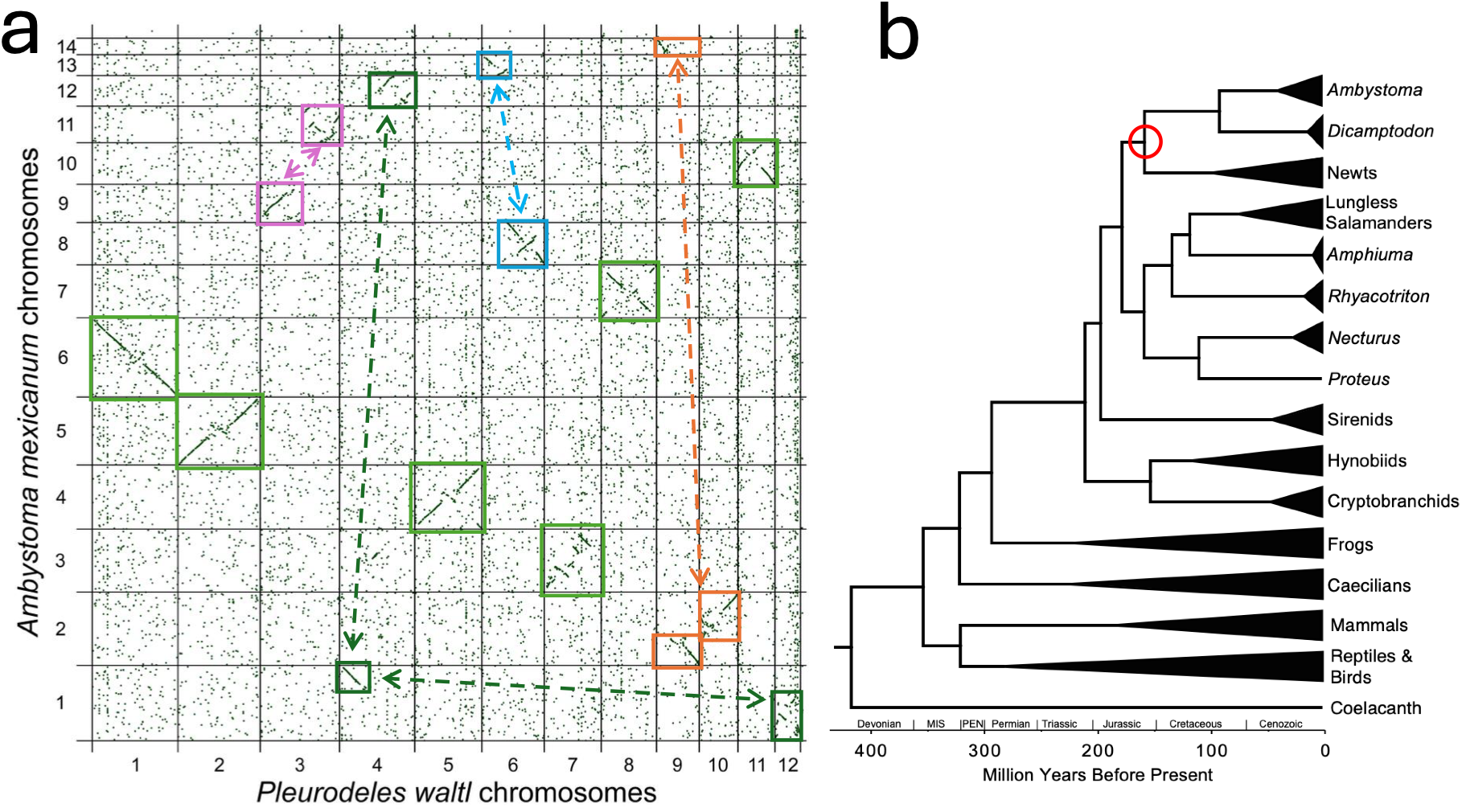
Comparison of axolotl and newt (*P. waltl*) genomes. **a)** A plot of showing the location of aligning sequences between axolotl and newt (*Pleurodeles waltl*) genomes. Diagonal features represent alignments between long colinear regions of chromosomes, with chromosomes varying in the number of inversions accumulated in these species over the last 159 million years. Light green squares show chromosomes with 1:1 orthology between species, and other colors (connected by arrows) show small groups of chromosomes that are interrelated by fission and fusion events. **b)** A phylogeny of tetrapod lineages, focusing on salamander diversity. Triangles represent larger groups of species that have been collapsed into a single group. The vertex of each triangle is positioned at the deepest divergence within the group. A red circle marks the evolutionary node spanned by comparisons in panel A.

**Figure 3:**
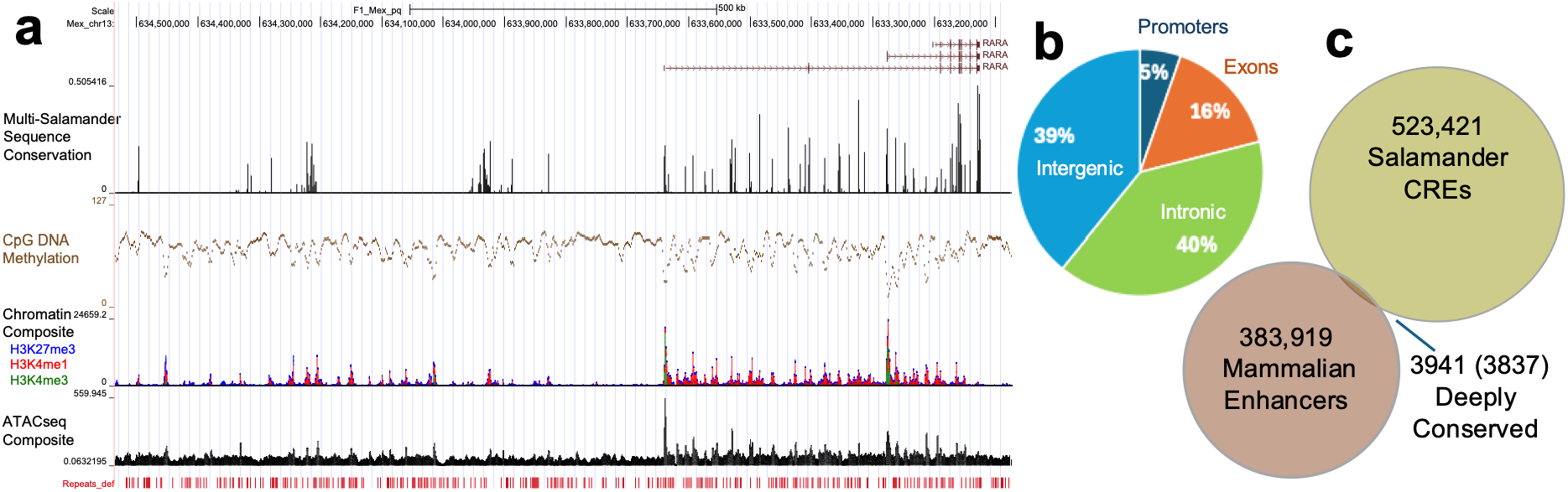
Identification of conserved elements in the axolotl genome. **a)** Analysis of sequence conservation identifies several peaks that visually overlap with published chromatin, ATAC and CpG methylation studies. The *RARA* (retinoic acid receptor alpha) gene body and upstream sequence is shown here. **b)** Nearly 80% of conserved sequences identified in multi-salamander alignments are present in intergenic and intronic regions. **c**) Overlap between identified salamander CREs and mammalian enhancer elements shows that only a small fraction of enhancers conserved in mammalian or salamander lineages are strongly conserved at the level of their primary DNA sequence. The two numbers reported for deeply conserved enhancers reflect the number of distinct enhancer elements in the human genome and the number of distinct conserved elements in axolotl (in parentheses).

These alignments identified 662,634 syntenic elements that are conserved between the axolotl and at least one other species that encompass coding sequences, proximate promoter regions, and conserved noncoding elements. In total we identified 523,421 intronic and intergenic elements ranging in size from 50bp to over 2kb (median length: 145bp, mean 313bp), that potentially represent conserved cis-regulatory elements (Figure 3).

### Assessment of conserved candidate cis-regulatory elements

We hypothesize that many of the conserved noncoding elements identified in our analyses correspond to cis-regulatory (enhancer-like) elements that are constrained due to their roles in regulating gene expression. As an initial test of this idea, we searched for a set of 383,919 nonredundant curated mammalian enhancers^38^ against the annotated axolotl genome. As would be expected given the observed high rates of turnover within mammalian lineages, we detected ∼1.2% of these in the axolotl genome. However, of the 4571 aligned mammalian elements 95% (4334) corresponded to 4206 conserved elements that were detected in multi-salamander alignments, the majority of which (3837) were classified as conserved intronic or intergenic sequences. These results show that the relatively small set of deeply conserved (ancestral tetrapod) cis-regulatory elements is well captured by the larger set of candidate enhancer elements that were independently identified in our multi-salamander alignments.

To more specifically test whether our salamander CREs reflect random matches between large genomes or sites that are likely to carry regulatory functions, we tested the degree to which these sequences overlap signals from previously-published epigenetic studies. To assess overlap between features and account for the differential influence of nucleosomes in shaping peak signals, we measured their co-occurrence in 300 bp bins across the genome. Conserved sequences occupied only 1.5% of the genome, whereas H3K4me1 and ATAC-seq peaks and methyl CpG-depleted regions occupied slightly larger fractions of the genome (8.6%, 8.8% and 7.8%) (Figure 3, Figure S2). The expected frequency of random overlap is extremely low (∼0.1%) given the large size of the genome (97 million bins across 29 Gb), or even the nonrepetitive fraction of the genome (9.6 Gb), however we found substantial overlap between all classes, with 23-32% of CREs overlapping with each chromatin-based enhancer set. This degree of overlap is notably lower than what is observed for conserved promoter peaks (Figure 4a,b), as might be expected given tissue specificity and higher rates of sequence turnover for enhancers ^20–22,38^. Although, the tissues sampled in these published epigenetic studies represent only a fraction of cell types and developmental states within the axolotl, we find clear orthogonal cis-regulatory signals for 76% of detected conserved elements, which can be further subdivided into definable enhancer states reflective of their average condition across regeneration studies (Figure 4c).

**Figure 4:**
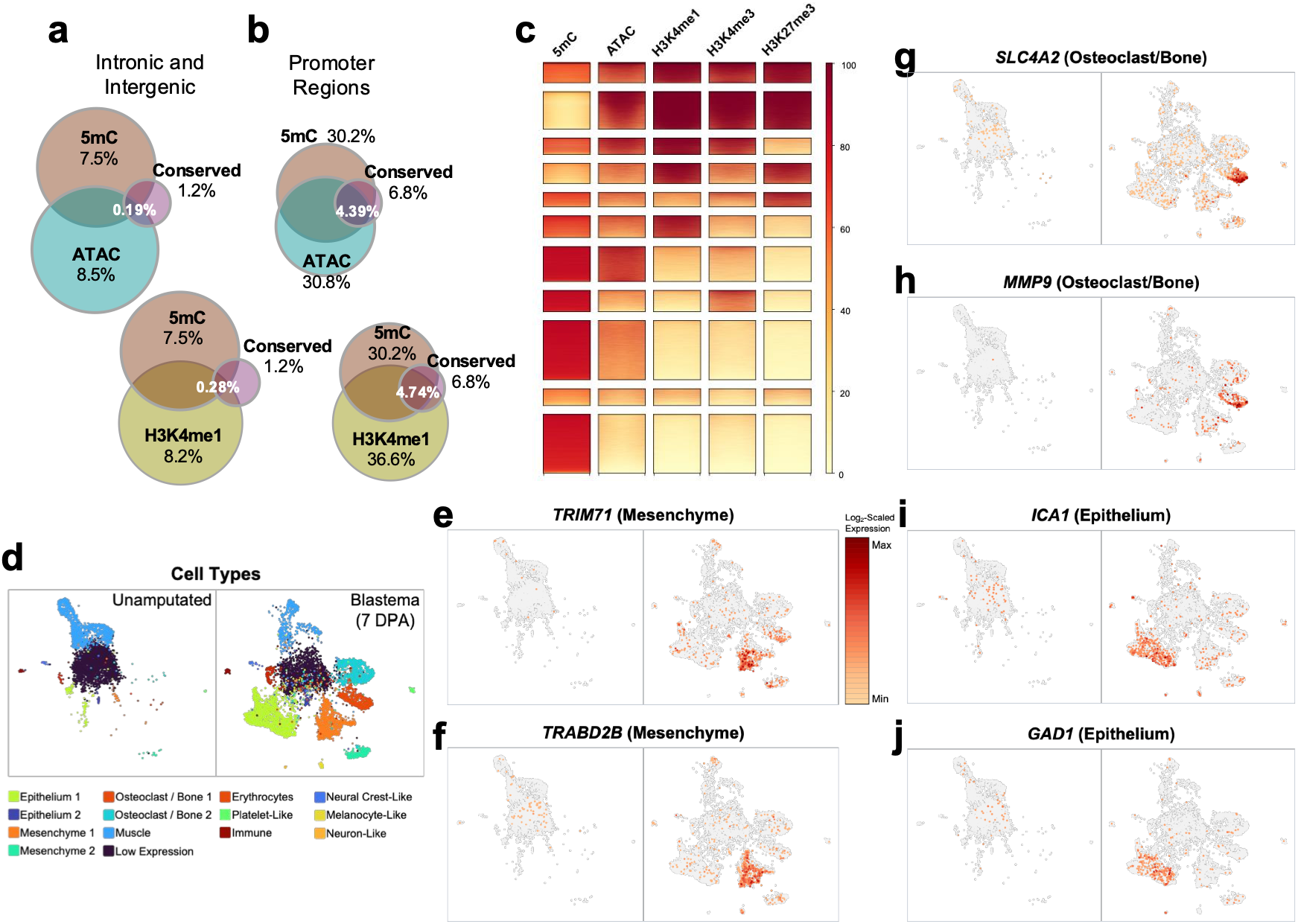
Salamander conserved elements identify a subset of cis-regulatory sequences, including sequences that are associated with upregulation of genes in specific cell types in the regeneration blastema. a&b) Overlap between salamander conserved sequences and epigenetic marks in **a)** intronic and intergenic noncoding regions, showing the percent of intergenic and intronic windows containing peaks for corresponding epigenetic marks and **b)** overlaps in promoter regions showing the percent of promoter windows containing peaks for corresponding epigenetic marks. **c)** Examining patterning of epigenetic marks across salamander conserved elements, averaged across diverse regenerating and non-regenerating tissues, shows that a small fraction of salamander conserved elements have been shown to be heterochromatic in samples studied to date, whereas the majority others show evidence of accessible chromatin or positive enhancer peaks. **d-j)** Assessing gene expression patterns associated with conserved regeneration enhancers using single cell sequencing of intact limb vs blastema (7 days post amputation). UMAP projections are based on combined expression and accessibility signals from both timepoints, with the two timepoints displayed separately (unamputated: left; blastema: right). **d)** Distinct cell clusters were identified based on shared nearest neighbor modularity optimization and are labeled based on their corresponding cell type. **e-j)** Expression of genes closest to several salamander conserved elements showing examples of cell type specific expression in mesenchyme (e,f), osteoclast/bone (g,h) and epithelium (i,j).

Studies of mammalian CREs have found that conserved elements generally have a larger influence on gene regulation than lineage-specific enhancers. As such, we reasoned that salamander-conserved CREs might play a stronger role during tissue regeneration, in comparison to nonconserved CREs. To test this expectation, we focused on epigenetic changes during limb blastema formation, comparing ATAC-seq data from intact limb to ATAC-seq data collected when progenitor cells begin to form a regeneration blastema at 3 days post amputation, and when blastemal cells begin to proliferate at 7 days post amputation (DPA)^39^. Focusing on genomic intervals that contained intronic and intergenic CREs that were defined from either ATAC or H3K4me1 datasets, we identified thousands of regions that change in chromatin accessibility during blastema formation (|log2(fold change)|>2). In total 1,750 ATAC-based and 5,045 H3K4me1-based CRE intervals were detected as differentially accessible during blastema initiation. A larger number (3,068 ATAC-based and 7,854 H3K4me1-based) were differentially accessible during early blastema proliferation. Comparing the fractions of conserved CREs vs nonconserved CREs that change in accessibility revealed that salamander-conserved CREs are 3-11X more likely to change when compared to either class of non-conserved CREs (Supplementary Table 2). These salamander-conserved CREs include the previously validated regeneration-specific axe.R enhancer, which is located in the first intron JAZF1 in the new assembly (Figure S4), and a total of 1507 distinct conserved blastema CREs (Supplementary Table 3). More broadly, differentially accessible conserved blastema CREs were found to occur near genes that are enriched in functions that are related to muscle skeletal and nervous system development, synapse assembly, extracellular matrix remodeling and cell migration, consistent with expected roles in regulating regeneration (Supplementary Table 4).

We expect that many of these conserved blastema CREs function as enhancers that regulate the expression of adjacent genes in specific cell types during blastema formation. To test this expectation, we performed single nucleus multiome (RNA-seq and ATAC-seq) analyses targeting 7 DPA forearm blastema and 0 DPA forearm stump tissue, sampling 7,712 and 3,913 cells respectively. These analyses captured all major cell types that participate in wound remodeling and blastema formation and allowed us to specifically examine the expression of genes that contained (for intronic CREs) or were nearest to the inferred set of conserved blastema CREs. These single cell analyses revealed that many of these presumptive enhancer target genes are upregulated in specific cell types during blastema formation, consistent with the prediction that many conserved blastema CREs encode precise enhancer-like activities during blastema formation (Figure 4, Supplementary Table 2).

In summary, our new axolotl assembly and multi-salamander alignments substantially extend the biological and genomic breadth of regulatory elements in salamander genomes, increase the interpretive value of epigenetic-based studies of regulatory elements in axolotl, and reveal the power of multispecies comparisons in identifying functional elements in even the largest vertebrate genomes. Notably, these alignments encompass only 5 of the 837 living salamander species, which share broadly archetypical tetrapod morphology, extensive life history variation and enhanced regenerative capacity. As high-quality reference genomes for additional large salamander genomes become available, we anticipate that extensions of similar synteny guided alignment approaches will further enrich our understanding of the function and evolution of regulatory sequences in these genomes.

## METHODS

### Animal Experimental Procedures

Animal procedures were approved by the University of Kentucky Institutional Animal Care and Use Committee (Protocol 2020-3449).

### *in vitro* Fertilization and Animal Rearing

F1 hybrids were generated by artificial insemination following previously published protocols^40^. The *A. mexicanum* female used in this cross was selected from the common laboratory stock maintained at the University of Kentucky Ambystoma Genetic Stock Center. This laboratory stock is the source of nearly all experimental animals used in modern studies and shares both ancestral and recently introgressed polymorphisms with incipient inbred strains ^18,19,26,27^. The *A. tigrinum* male was collected from a wild population in Kentucky, in a pond maintained by the Clay Hill Memorial Forest (Campbellsville University, Campbellsville, KY), under Kentucky Collection permit SC2311123.

### Sequencing and Assembly

High molecular weight DNA was extracted from a single female interspecific F1 hybrid using the Monarch® HMW DNA Extraction Kit for Cells & Blood (New England Biolabs). This DNA was used for library preparation and HiFi sequencing on 18 PacBio SMRT cells (24 hour movie time) in a PacBio Revio sequencer by the HudsonAlpha Genome Sequencing Center. DNA from the *A. mexicanum* and *A. tigrinum* parents of this animal was extracted by standard phenol/chloroform isolation. Illumina library construction and sequencing of parental DNAs (2×150 bp reads) was performed using the Illumina DNA PCR-Free Prep kit and an Illumina NovaSeq S4 by the Roy J. Carver Biotechnology Center University of Illinois at Urbana-Champaign. Arima Hi-C library construction used the regenerated tail tip (approximately one month post amputation) of the F1 hybrid. Armia Hi-C libraries were prepared for this sample using published VGP protocols^41^, and sequenced on an Illumina NovaSeq S4 by the Vertebrate Genomics Laboratory (The Rockefeller University).

To aid in haplotype resolution kmer hash tables were first built for both parental species using yak-0.1 (https://github.com/lh3/yak) and used alongside PacBio HiFi reads to generate a trio-binned assembly with Hifiasm-0.19.6-r595^42–44^. Initial attempts to integrate Hi-C reads during Hifiasm assembly resulted in recurrent segfaults, necessitating later integration of Hi-C datasets. After generation of species-phased contigs via Hifiasm, Hi-C reads were mapped to the genome and filtered using the VGP protocol^15,45^. Hi-C reads were then used to scaffold the assembly using yahs^46^. Following initial scaffolding, Hi-C data were remapped to both genome haplotype assemblies using BWA^42^ to permit manual curation using PretextView (https://github.com/sanger-tol/PretextView). This necessitated splitting several of the longer scaffolds to a length of less than 2.1Gb at scaffold breaks to account for the use of 32-bit integers in mapping algorithms. Following manual assembly curation, larger contigs were split at scaffolding gaps within centromeric regions to permit ingestion into NCBI and other databases that rely on 32-bit indexing.

### Methylation Analysis

CpG methylation tracks were generated from the same PacBio HiFi reads that were used to generate the assembly. Reads were aligned to the genome using pbmm2 align (v.1.16.99: https://github.com/PacificBiosciences/pbmm2), CpG scores were calculated using pb-CpG-tools-v2.3.2 (https://github.com/PacificBiosciences/pb-CpG-tools). Dips in CpG methylation (indicative regions of open chromatin) were identified by calling peaks using MACS3 (v3.0.0a7)^47,48^ bdgpeakcall with cutoff 70, minimum length 100 and maximum gap 200 (-c 70, -l 100, -g 200) on inverted methylation scores.

### Analysis of ChIPseq and ATAC-seq datasets

We downloaded previously published ATAC-seq (PRJNA682840)^39^ and CUT&Tag datasets (H3K4me1, H3K4me3 and H3K27me3: PRJNA899786)^49^ from regeneration blastemas and amputated limbs. ATAC-seq and CUT&Tag reads were trimmed using Trim Galore (v.0.6.10: https://github.com/FelixKrueger/TrimGalore) and aligned with bwa mem (v.0.7.18)^50^. Then, after removing duplicated, secondary, auxiliary alignments, and alignments with more than 5 mismatches, RPM-normalized read coverage was calculated across all genomic bases. ATAC-seq and H3K4me1peaks were called for individual samples using MACS3 (v3.0.0a7)^47,48^ bdgpeakcall command with parameters minimum length of peak -l 100, maximum gap -g 200, and empirically defined cutoff values of -c 0.009 for ATAC-seq coverage and -c 0.06 for H3K4me1. Peaks called for individual samples were merged across all samples to produce unified sets of H3K4me1-based and ATAC-based peaks. To account for differential positioning relative to nucleosomes and uniformly assess overlap between conservation sites and peaks called for ATAC-seq, 5mC and H3K4me1, we measured their co-occurrence in 300 bp bins across the genome using BEDtools intersect -c (BEDtools v2.27.1)^51^.

We used deepTools v3.5.6^52^ to calculate and plot profiles across averaged gene and CRE features. First, we used bigwigAverage with a default bin size 50bs to calculate average coverage across all samples for each chromatin mark. Then the computeMatrix reference-point tool was used to calculate average coverage depth profiles for each chromatin mark and average methylation scores profiles within a 3 Kb distance from transcription start sites (TSS) of 24,056 annotated primary mRNAs (coding gene isoforms: Supplementary Table 5).

For metagene analyses of ATAC-seq enrichment we applied a computeMatrix scale-regions tool with options --metagene, -a 500, -b 500, -m 1000, --binSize 10 to calculate a metagene coverage profile for the same annotated primary mRNAs. To perform metagene analyses of methylation patterns in the axolotl we first calculated average methylation scores for 1 kb windows with a 10 bs sliding step, and then followed the same profiling method used for ATAC-seq. For the human metagene analyses, we used MANE gene annotation v1.4. Methylation scores, defined as percentage of reads methylated at each CpG site, for the human genome were obtained from methbase (https://github.com/smithlabcode/methbase)^53^ and were derived from SRA records SRX27441857 and SRX27441858. Finally, we applied the same strategy used for *A. mexicanum* to generate methylation profiles across human metagenes.

### Clustering of Chromatin States Across Salamander-Conserved Elements

To characterize conserved elements located in introns or intergenic regions, we considered 500bp intervals centered on small elements and wider intervals centered on larger elements (251-2000 bs) corresponding to 2X the element’s length. For methylation, we utilized the same bigwig that was generated for metagene analysis. For other chromatin marks, to avoid the effect of outliers, all values above the M=90^th^ percentile were replaced with M, and all values below m=10^th^ percentile were replaced with m. All values were then rescaled to a [0,100] range to match the methylation scores range. Finally, computeMatrix scale-regions with --regionBodyLength 500 and plotHeatmap with k-means clustering algorithm (k=11) were used to identify similarly marked groups of elements and generate heatmaps.

### Alignment of Enhancers from Divergent Mammalian Lineages

We downloaded a curated list of hg38 intervals for putative regulatory elements conserved across mammalian lineages^38^, merged overlapping intervals using bedtools merge (BEDtools v2.27.1)^51^ and extracted 383,919 corresponding sequences from hg38. These sequences were aligned to the axolotl genome using BLASTN v.2.11.0^54,55^ with parameter -dust set to “no”. Alignments were filtered to retain only hits whose length was either greater than half of the query length or exceeding 99 bases. In addition to all unique alignments, for elements aligned to multiple locations alignment with a maximum bitscore was added to the set of primary alignments. Other alignments formed a secondary group. Finally, overlapping alignment intervals on the axolotl genome were merged across gaps of less than or equal to 50 base pairs.

### Single Cell Multiome Processing and Analysis

Eight juvenile axolotls (RRID:AGSC_101S; 298-305 days post fertilization) were obtained from the Ambystoma Genetic Stock Center (AGSC RRID: SCR_006372) and administered midpoint forelimb amputations under 0.02% benzocaine anesthesia. At the time of amputation and 7 days post-amputation, axolotls were administered benzocaine anesthesia and 1.5 mm of tissue was collected from distal forelimb stumps to generate three independent samples (four limb stumps each). The samples were shipped on dry ice to Singulomics, where nuclei were isolated for each sample and then all nuclei were pooled to generate sequence libraries using the 10x Next GEM Single Cell Multiome ATAC + Gene Expression kit. Each library was then sequenced on an Illumina NovaSeq 6000 to generate with ∼200 million 150 bp pair end reads. Cell Ranger ARC (v.2.0.2) was used to perform initial analysis of the resulting sequence sets. To meet software constraints, chromosome-size scaffolds in the reference genome were split into two to four chunks, ensuring that no chunk exceeded 536.8 Mb. Split coordinates for each chromosome were selected so that the divisions occurred within scaffold gaps and did not disrupt any genes (the corresponding AGP file is uploaded to the GOE record for this study: GSE343762). Human homolog names were manually assigned to 8,285 genes annotated but lacked names in the RefSeq annotation (Supplementary Table 6). We used AGAT (v.1.6.1) to convert gene annotation from GFF to GTF, and the genes’ coordinates were lifted over to the split reference via CrossMap.py (v.0.3.6). Only protein-coding genes and long non-coding RNAs were used to prepare the reference with cellranger-arc mkref. The two datasets were processed using cellranger-arc count, and the results were aggregated via cellranger-arc aggr. This aggregated dataset was then reprocessed using Seurat to generate a UMAP projection and define cell clusters based on multimodal weighted nearest neighbor analysis using the function FindMultiModalNeighbors^56,57^. Ontology enrichment analysis was performed using the ENRICHR server^58^.

## Data Availability

The axolotl genome assembly and genome sequence datasets are released under bioproject PRJNA1104202 and under the VGP umbrella project PRJNA489243. Sequencing reads for F1 *A. mexicanum* X *A. tigrinum* hybrid (NCBI SRA accession number SRR40236586) are also placed under the bioproject PRJNA1104202. The *A. tigrinum* genome assembly and genome sequence datasets are released under bioproject PRJNA1512557. The bioprojects PRJNA1104202 and PRJNA1512557 are both linked to umbrella project PRJNA1168523. Single cell sequence data and analyses are deposited in the NCBI GEO repository (accession number GSE343726).

## Acknowledgements

This work was supported by the National Institutes of Health (OD010435 to S.R.V. and J.J.S; OD019794 to S.R.V). We thank Amy Berry (Clay Hill Memorial Forest, Campbellsville University) for assistance in obtaining the tiger salamander used for this study. The authors acknowledge the support of the Ambystoma Genetic Stock Center (AGSC_RRID:SCR_006372), the University of Kentucky High-Performance Computing complex, and the Oncogenomics Shared Resource of the University of Kentucky Markey Cancer Center which is supported by the National Institutes of Health (P30CA177558).

## References

1. Kardong, K.V. Vertebrates : comparative anatomy, function, evolution, xix, 790 pages (McGraw-Hill Education, New York, NY, 2019).

2. Kang, W., Fei, J.F., Liang, C., Wagner, G. & Zhou, Q. Evolution of limb and digit identity genes since the tetrapod ancestor. Nat Commun 17(2026).

3. Onimaru, K. et al. Developmental hourglass and heterochronic shifts in fin and limb development. Elife 10(2021).

4. Shubin, N., Tabin, C. & Carroll, S. Deep homology and the origins of evolutionary novelty. Nature 457, 818–23 (2009).

5. Woltering, J.M. & Duboule, D. Tetrapod axial evolution and developmental constraints; Empirical underpinning by a mouse model. Mech Dev 138 Pt 2, 64–72 (2015).

6. Johnson, C.K. & Voss, S.R. Salamander paedomorphosis: linking thyroid hormone to life history and life cycle evolution. Curr Top Dev Biol 103, 229–58 (2013).

7. Jia, J. et al. Ossification patterns of the carpus and tarsus in salamanders and impacts of preaxial dominance on the fin-to-limb transition. Sci Adv 8, eabq7669 (2022).

8. Trofka, A. et al. Genetic basis for an evolutionary shift from ancestral preaxial to postaxial limb polarity in non-urodele vertebrates. Curr Biol 31, 4923–4934 e5 (2021).

9. Joven, A., Elewa, A. & Simon, A. Model systems for regeneration: salamanders. Development 146(2019).

10. Sessions, S.K. Evolutionary cytogenetics in salamanders. Chromosome Res 16, 183–201 (2008).

11. Sessions, S.K. & Wake, D.B. Forever young: Linking regeneration and genome size in salamanders. Dev Dyn 250, 768–778 (2021).

12. Stewart, A.A. & Wiens, J.J. A time-calibrated salamander phylogeny including 765 species and 503 genes. Mol Phylogenet Evol 204, 108272 (2025).

13. Smith, J.J. et al. Genic regions of a large salamander genome contain long introns and novel genes. BMC Genomics 10, 19 (2009).

14. Keinath, M.C. et al. Initial characterization of the large genome of the salamander Ambystoma mexicanum using shotgun and laser capture chromosome sequencing. Sci Rep 5, 16413 (2015).

15. Formenti, G., et al. The Vertebrate Genomes Project Phase I: A global reference genome resource. bioRxiv, 2026.06.24.732306 (2026).

16. Nurk, S. et al. The complete sequence of a human genome. Science 376, 44–53 (2022).

17. Pruitt, K.D. et al. RefSeq: an update on mammalian reference sequences. Nucleic Acids Res 42, D756–63 (2014).

18. Schloissnig, S. et al. The giant axolotl genome uncovers the evolution, scaling, and transcriptional control of complex gene loci. Proc Natl Acad Sci U S A 118(2021).

19. Smith, J.J. et al. A chromosome-scale assembly of the axolotl genome. Genome Res 29, 317–324 (2019).

20. Berthelot, C., Villar, D., Horvath, J.E., Odom, D.T. & Flicek, P. Complexity and conservation of regulatory landscapes underlie evolutionary resilience of mammalian gene expression. Nat Ecol Evol 2, 152–163 (2018).

21. Christmas, M.J. et al. Evolutionary constraint and innovation across hundreds of placental mammals. Science 380, eabn3943 (2023).

22. Dukler, N., Huang, Y.F. & Siepel, A. Phylogenetic Modeling of Regulatory Element Turnover Based on Epigenomic Data. Mol Biol Evol 37, 2137–2152 (2020).

23. Foley, N.M. et al. A genomic timescale for placental mammal evolution. Science 380, eabl8189 (2023).

24. Phan, M.H.Q., et al. Conservation of regulatory elements with highly diverged sequences across large evolutionary distances. Nat Genet 57, 1524–1534 (2025).

25. Armstrong, J. et al. Progressive Cactus is a multiple-genome aligner for the thousand-genome era. Nature 587, 246–251 (2020).

26. Nowoshilow, S. et al. The axolotl genome and the evolution of key tissue formation regulators. Nature 554, 50–55 (2018).

27. Woodcock, M.R. et al. Identification of Mutant Genes and Introgressed Tiger Salamander DNA in the Laboratory Axolotl, Ambystoma mexicanum. Sci Rep 7, 6 (2017).

28. Timoshevskaya, N., Voss, S.R., Labianca, C.N., High, C.R. & Smith, J.J. Large-scale variation in single nucleotide polymorphism density within the laboratory axolotl (Ambystoma mexicanum). Dev Dyn 250, 822–837 (2021).

29. Pacheco-Olvera, D.L. et al. Genomic characterization of the T-cell receptor loci in Ambystoma mexicanum. Front Immunol 16, 1656386 (2025).

30. Thibaud-Nissen, F., Souvorov, A., Murphy, T., DiCuccio, M. & Kitts, P. Eukaryotic Genome Annotation Pipeline. in The NCBI Handbook [Internet]. Available from: https://www.ncbi.nlm.nih.gov/books/NBK1CS43S/ (ed. (US), N.C.f.B.I.) (Bethesda (MD), 2013).

31. Cooper, G.M. et al. Distribution and intensity of constraint in mammalian genomic sequence. Genome Res 15, 901–13 (2005).

32. Felsenstein, J. & Churchill, G.A. A Hidden Markov Model approach to variation among sites in rate of evolution. Mol Biol Evol 13, 93–104 (1996).

33. Pollard, K.S., Hubisz, M.J., Rosenbloom, K.R. & Siepel, A. Detection of nonneutral substitution rates on mammalian phylogenies. Genome Res 20, 110–21 (2010).

34. Siepel, A. et al. Evolutionarily conserved elements in vertebrate, insect, worm, and yeast genomes. Genome Res 15, 1034–50 (2005).

35. Streicher, J.W. et al. The genome sequence of the great crested newt, Triturus cristatus (Laurenti, 1768). Wellcome Open Res 10, 270 (2025).

36. Keinath, M.C., Voss, S.R., Tsonis, P.A. & Smith, J.J. A linkage map for the Newt Notophthalmus viridescens: Insights in vertebrate genome and chromosome evolution. Dev Biol 426, 211–218 (2017).

37. Chandra, G., Vasimuddin, M., Misra, S. & Jain, C. Accelerating minimap2 for whole-genome alignment. Bioinformatics 42(2026).

38. Uebbing, S. et al. Evolutionary Innovations in Conserved Regulatory Elements Associate With Developmental Genes in Mammals. Mol Biol Evol 41(2024).

39. Wei, X. et al. An ATAC-seq Dataset Uncovers the Regulatory Landscape During Axolotl Limb Regeneration. Front Cell Dev Biol 9, 651145 (2021).

40. Armstrong, J.B. & Duhon, S.T. Induced spawnings, artificial insemination, and other genetic manipulations. in Developmental Biology of the Axolotl (eds. Armstrong, J.B. & Malacinski, G.) 228–236 (Oxford University Press, New York, NY, 1989).

41. Rhie, A. et al. Towards complete and error-free genome assemblies of all vertebrate species. Nature 592, 737–746 (2021).

42. Cheng, H., Asri, M., Lucas, J., Koren, S. & Li, H. Scalable telomere-to-telomere assembly for diploid and polyploid genomes with double graph. Nat Methods 21, 967–970 (2024).

43. Cheng, H., Concepcion, G.T., Feng, X., Zhang, H. & Li, H. Haplotype-resolved de novo assembly using phased assembly graphs with hifiasm. Nat Methods 18, 170–175 (2021).

44. Cheng, H. et al. Haplotype-resolved assembly of diploid genomes without parental data. Nat Biotechnol 40, 1332–1335 (2022).

45. Lariviere, D. et al. Scalable, accessible and reproducible reference genome assembly and evaluation in Galaxy. Nat Biotechnol 42, 367–370 (2024).

46. Zhou, C., McCarthy, S.A. & Durbin, R. YaHS: yet another Hi-C scaffolding tool. Bioinformatics 39(2023).

47. Feng, J., Liu, T., Qin, B., Zhang, Y. & Liu, X.S. Identifying ChIP-seq enrichment using MACS. Nat Protoc 7, 1728–40 (2012).

48. Zhang, Y. et al. Model-based analysis of ChIP-Seq (MACS). Genome Biol 9, R137 (2008).

49. Kawaguchi, A. et al. A chromatin code for limb segment identity in axolotl limb regeneration. Dev Cell 59, 2239–2253 e9 (2024).

50. Li, H. & Durbin, R. Fast and accurate short read alignment with Burrows-Wheeler transform. Bioinformatics 25, 1754–60 (2009).

51. Quinlan, A.R. & Hall, I.M. BEDTools: a flexible suite of utilities for comparing genomic features. Bioinformatics 26, 841–2 (2010).

52. Ramirez, F. et al. deepTools2: a next generation web server for deep-sequencing data analysis. Nucleic Acids Res 44, W160–5 (2016).

53. Song, Q., et al. A reference methylome database and analysis pipeline to facilitate integrative and comparative epigenomics. PLoS One 8, e81148 (2013).

54. Altschul, S.F., Gish, W., Miller, W., Myers, E.W. & Lipman, D.J. Basic local alignment search tool. J Mol Biol 215, 403–10 (1990).

55. Camacho, C. et al. BLAST+: architecture and applications. BMC Bioinformatics 10, 421 (2009).

56. Hao, Y. et al. Integrated analysis of multimodal single-cell data. Cell 184, 3573–3587 e29 (2021).

57. Hao, Y. et al. Dictionary learning for integrative, multimodal and scalable single-cell analysis. Nat Biotechnol 42, 293–304 (2024).

58. Chen, E.Y. et al. Enrichr: interactive and collaborative HTML5 gene list enrichment analysis tool. BMC Bioinformatics 14, 128 (2013).

